# Global changes in alternative splicing are associated with insecticide response and resistance

**DOI:** 10.1101/2025.10.22.684019

**Authors:** Christine A. Tabuloc, Sergio Hidalgo, Curtis R. Carlson, Hongtao Zhang, Frank G. Zalom, Joanna C. Chiu

## Abstract

Alternative splicing (AS) promotes phenotypic plasticity to adverse conditions by altering transcripts involved in stress adaptation. However, whether changes in AS occur transcriptome-wide or in only a few key genes is unclear. Agricultural insect pests are regularly exposed to xenobiotic stress from insecticide applications and thus develop resistance. We show that the pest *Drosophila suzukii,* resistant to multiple insecticides, exhibits increased global AS events compared to susceptible flies. Alternatively spliced genes are enriched in multiple processes including stress response and insecticide resistance, suggesting AS is a mechanism underlying insecticide resistance development. Furthermore, sublethal insecticide exposure promotes AS events even in the absence of substantial differential gene expression. This study provides insights into the role of AS in enabling insects to diversify genome function to survive acute insecticide treatment and to develop xenobiotic resistance.

## Introduction

In nature, organisms experience diverse changes in their environments, such as extreme weather conditions, fluctuations in nutrient availability, and exposure to predators, toxins, and xenobiotics. In order to survive these challenges, organisms need to adapt quickly and continuously. There are many known molecular mechanisms responsible for driving adaptations. One example is alternative splicing (AS) (reviewed in (Singh and Ahi 2022; Verta and Jacobs 2022)). AS is the process that produces different RNA isoforms encoded from the same gene, which can result in, for instance, different proteins varying in structure and function or even producing transcripts targeted for degradation prior to protein translation (reviewed in (Nilsen and Graveley 2010)). In both plants and animals, AS is important for enabling phenotypic plasticity and adaptation to many abiotic stresses, including temperature variations (Lee et al. 2013; Posé et al. 2013; Horii et al. 2019; Martin Anduaga et al. 2019), pathogen infection (Yukl et al. 2018; Chauhan et al. 2019; Annalora et al. 2024), nutrient stress (reviewed in (Laloum et al. 2018)), and insecticide exposure (Shao et al. 2007; Dong et al. 2014; Berger et al. 2016; Wu et al. 2017; Ureña et al. 2019). However, many of these studies focus on a few genes previously characterized to be necessary in these processes or relied on technology that could not accurately measure all the expressed isoforms.

Considering that AS is important in responding to diverse environmental stressors, it is possible that changes in AS may not be limited to a few genes but rather occur transcriptome-wide. Consequently, under stressful conditions, only some genes that are differentially spliced promote adaptations and increase survival, while others might not be involved in this response at all and could even incur fitness costs. Recent advancements in long-read sequencing now enable the survey of the entire AS landscape with increased precision (Xie et al. 2020; Kanwar et al. 2021), enabling the testing of this hypothesis. Here, we sought to investigate whether changes in AS occur transcriptome-wide to drive insecticide resistance in *Drosophila suzukii*, a worldwide invasive pest of fruit crops heavily targeted by insecticide applications.

## Results

### Insecticide-resistant flies exhibit an altered alternative splicing landscape compared to susceptible lines

Insecticide resistance is a significant challenge in agriculture. It has been associated with target-site resistance, metabolic resistance, penetration resistance and even behavioral resistance (reviewed in (Siddiqui et al. 2023)). Several genes encoding proteins targeted by insecticides are differentially spliced, and this aberrant splicing has been linked to resistance (Hemingway et al. 1998; Grauso et al. 2002; Shao et al. 2007; He et al. 2012; Berger et al. 2016; Wu et al. 2017; Ureña et al. 2019; Shen et al. 2021). We hypothesize that alternative splicing (AS) patterns could be altered transcriptome-wide in insecticide-resistant flies rather than only affecting one or a few target genes. To test our hypothesis, we first sought to accurately account for the AS events in *Drosophila suzukii.* Leveraging a combination of short- and long-read sequencing, we generated a transcriptome re-annotation with improved isoform discovery (**S. Tables 1-3**). With this annotation, we captured the AS landscape in two isofemale resistant lines to Entrust (Entrust_Resis1 or Entrust_Resis2), a spinosyn-type insecticide commonly used to manage *D. suzukii*, by comparing their isoform-informed transcriptome with the one generated from two pooled isofemale susceptible lines (Entrust_Sus1 and Entrust_Sus2) developed from a single population of field-collected flies from California, USA (**Fig. 1; S. Tables 4-6**). Pooling the susceptible lines accounts for biological differences not attributed to resistance (Cridland et al. 2023; Tabuloc et al. 2024). We then obtained a baseline AS landscape by comparing the two susceptible lines (i.e. Entrust_Sus1 versus Entrust_Sus2) and compared the AS events that arise from this comparison to the AS events obtained from each resistant line compared to the pooled susceptible lines. We observed an increase in differential splicing events in the Entrust_Resis2 line as compared to the AS events of the Entrust-susceptible baseline (Fisher’s Exact Test=2.62e-14, q=5.24e-14), while there was no difference in AS events in Entrust_Resis1 line as compared to the Entrust-susceptible baseline (Fisher’s Exact Test=0.7415, q=0.7415; **Fig. 1A**). Increases in AS events were also observed for the lines resistant to Mustang Maxx (i.e. MustangMaxx_Resis1 or MustangMaxx_Resis2 versus pooled Mustang Maxx-susceptible lines), a pyrethroid-type insecticide also used to manage *D. suzukii*, as compared to the susceptible control comparison (MustangMaxx_Sus1 versus MustangMaxx_Sus2) (MustangMaxx_Resis1 Fisher’s Exact Test=5.63e-05, q=1.1264e-4; MustangMaxx_Resis2 Fisher’s Exact Test=5.68e-4, q=5.68e-4) (**Fig. 1B; S. Tables 7-9**). Because Entrust_Resis1 exhibits penetration resistance, which is an upregulation of cuticular genes, and Entrust_Resis2, MustangMaxx_Resis1, and MustangMaxx_Resis2 all exhibit metabolic resistance, which is an upregulation of metabolic detoxification enzymes (Tabuloc et al. 2024), our observations suggest a possible association between metabolic resistance and changes in AS. We then determined how many genes exhibited at least one AS event (**Fig. 1C-D**). Similar to what we observed with the occurrence of AS events, we determined that lines exhibiting metabolic resistance (Entrust_Resis2 and both Mustang Maxx-resistant lines) have more differentially spliced genes compared to the susceptible line comparisons than the comparison with the line exhibiting penetration resistance (Entrust_Resis1 Fisher’s Exact Test=0.5261, q=0.5261; Entrust_Resis2 Fisher’s Exact Test=4.08e-6, q=8.16e-06; MustangMaxx_Resis1 Fisher’s Exact Test=0.009189, q=0.018378; MustangMaxx_Resis2 Fisher’s Exact Test=0.02334, q=0.02334).

**Figure 1:**
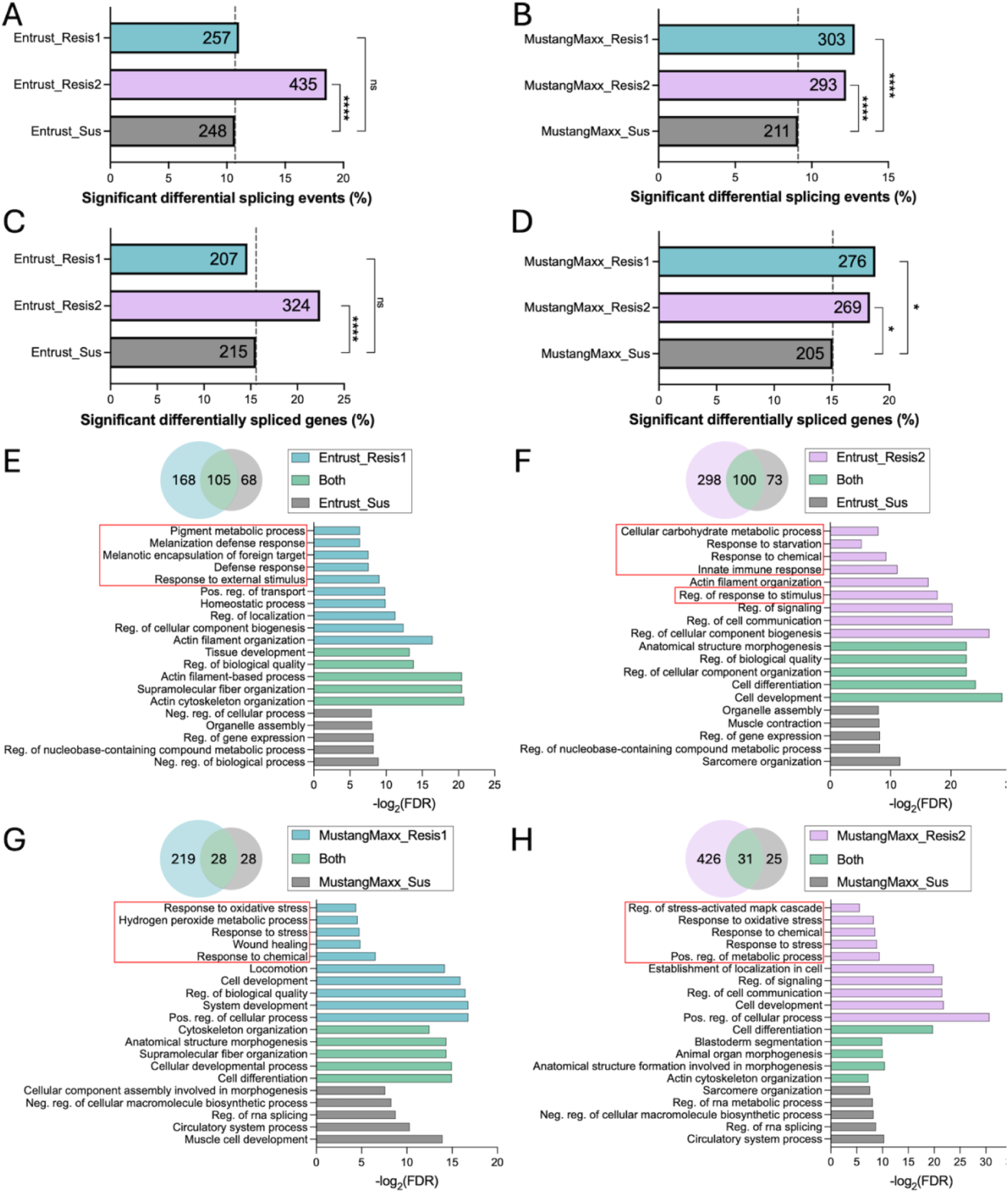
Entrust- and Mustang Maxx-resistance is associated with increased alternative splicing. (A-B) Percentage of significant differential alternative splicing (AS) events out of all detected AS events in either (A) Entrust-resistant lines (Entrust_Resis1 (blue) and Entrust_Resis2 (pink)) vs Entrust-susceptible lines or (B) Mustang Maxx-resistant lines (MustangMaxx_Resis1 (blue) and MustangMaxx_Resis2 (pink)) vs the Mustang Maxx-susceptible lines. The number of significant AS events is displayed within each bar. The dotted line indicates the % of significant differential splicing events, defined as the baseline AS landscape, in the susceptible groups. (C-D) Percentage of significant differentially spliced genes of all detected alternatively spliced genes in either (C) Entrust-resistant lines (Entrust_Resis1 (blue) and Entrust_Resis2 (pink)) vs the Entrust-susceptible lines or (D) Mustang Maxx-resistant lines (MustangMaxx_Resis1 (blue) and MustangMaxx_Resis2 (pink)) vs the Mustang Maxx-susceptible lines. The dotted line marks the % of significant differentially spliced genes in the susceptible group. Asterisks denote significant p-values as determined by Fisher’s Exact test, corrected with Benjamini and Hochberg: * p > 0.05 and **** p < 0.0001. (E-H) Top: Venn diagrams summarizing the number of processes within the Gene Ontology (GO) Biological Processes categories that are enriched in the differentially spliced genes. Bottom: The bar graphs contain the top 5 most significant GO categories that are in the resistant line alone ((E,G) blue or (F,H) pink), the susceptible line alone (grey), and shared categories between the resistant and susceptible lines (green) as well as categories that are related to stress response processes or processes previously implicated in resistance (red box), such as (E) cuticle-related processes or (F-H) metabolism-related processes. The x-axis is the -log_2_(FDR), where FDR is the false discovery rate correction of enrichment p-values. For categories that are in both the resistant and susceptible lines, only the -log_2_(FDR) of the resistant line is shown.

We then performed enrichment analysis to determine Gene Ontology (GO) Biological Processes classifications for the differentially spliced genes (**Fig. 1E-H; S. Tables 10-15**). We observed that the AS genes in the Entrust_Resis1 comparison are enriched in a total of 273 processes, with 168 enriched only in the Entrust_Resis1 comparison and not in the Entrust-susceptible group comparison (**Fig. 1E; S. Table 10**). Interestingly, although there was no change in the number of AS events nor the number of AS genes for Entrust_Resis1, which has been shown to exhibit penetration resistance (Tabuloc et al. 2024) (**Fig. 1A and C**), we observed that the AS genes are enriched in processes pertaining to the insect cuticle, such as *melanization defense response* and *melanotic encapsulation of foreign target*, as well as environmental sensing such as *response to external stimulus*. This suggests that although no difference is observed in the number of events or genes, there is still differential splicing occurring in specific genes that could contribute to resistance. Overlapping processes between the Entrust_Resis1 comparison and the susceptible baseline comparisons include those related to the cytoskeleton, such as *actin cytoskeleton organization* while in the susceptible baseline comparisons, the most significant are housekeeping processes including *organelle assembly* and *regulation of gene expression*. When comparing Entrust_Resis2 to susceptible lines, AS genes were enriched in a total of 398 biological processes, with 100 of those GO terms also enriched in the susceptible baseline comparison (**Fig. 1F; S. Table 12**). Notably, AS genes are enriched in processes related to metabolism, such as *cellular carbohydrate metabolic process*, and stress response, such as *response to chemical* and *innate immune response*, which aligns with the observation that Entrust_Resis2 exhibits metabolic resistance (Tabuloc et al. 2024). Consistent with this observation, MustangMaxx_Resis1 and MustangMaxx_Resis2, both metabolically resistant lines, exhibit AS in genes enriched in processes related to stress response (e.g. *response to stress* and *response to chemical*) and metabolism (e.g. *hydrogen peroxide metabolic process* and *positive regulation of metabolic process*) (**Fig. 1G-H; S. Table 13-15**).

We next determined whether a particular AS type could be associated with resistance (**S. Fig. 1**). We categorized the splicing events into 7 groups: exon skipping (SE), intron retention (RI), mutually exclusive exon (MX), alternative first exon (AF), alternative last exon (AL), alternative 5’ splice site (A5), and alternative 3’ splice site (A3) (**S. Fig. 1A**) and assessed both the total number of AS events and the number of genes that exhibited each AS type. Overall, we found no evidence suggesting that a particular AS type accounts for the splicing changes observed in the resistant line comparisons (**S. Fig.1 B-E**). We observed a small, albeit significant, decrease in RI events in the Entrust_Resis2 comparison (AS events: Fisher’s Exact Test=0.0027, q=0.0189; AS genes: Fisher’s Exact Test=0.002478, q=0.019236) (**S. Fig. 1B and D**). Although there was no particular AS type favored overall in genes that were alternatively spliced, we could not yet rule out that a certain AS type is associated with genes previously reported to be involved in insecticide resistance. Thus, we looked specifically at genes involved in either metabolic (for the Entrust_Resis2, MustangMaxx_Resis1, and MustangMaxx_Resis2 comparisons) or cuticular processes (for the Entrust_Resis1 comparison). We only observed more AF events and fewer RI events in the Entrust_Resis2 comparison as compared to the pooled susceptible lines comparison (AF: Fisher’s Exact Test=0.0004148, q=0.0029036; RI: Fisher’s Exact Test=0.0008457, q=0.00295995) (**S. Fig. 1F**).

Our analyses of *Drosophila suzukii* flies resistant to two different commonly used insecticides suggest that lines exhibiting metabolic resistance undergo more AS events than the susceptible lines. Although no changes were observed in the number of AS in the line exhibiting penetration resistance, there were changes with respect to the functional enrichment of the differentially spliced genes. Overall, we observed that differential splicing appears to be widespread in all resistant lines, affecting many classes of genes involved in a range of processes. Some of these processes include those related to stress response or were previously described as contributing to insecticide resistance mechanisms such as metabolic and cuticular processes.

### Metabolic resistance is associated with aberrant splicing

To validate the association between global aberrant splicing and insecticide resistance, we analyzed *D. suzukii* flies resistant to the organophosphate Malathion, a third class of insecticide used to manage this pest. We first established new isofemale lines from a field-collected population exhibiting resistance to Malathion. Using discriminating dose bioassays (Gress and Zalom 2022), we assessed the resistance status of multiple isofemale lines (**Fig. 2A**) and identified that lines Malathion_Resis1 and Malathion_Resis2 are more resistant than a known susceptible control line also collected in California, USA, herein referred to as Wolfskill (Gress and Zalom 2019) (Wolfskill vs Malathion_Resis1: adjusted p<0.0001, t=13.57, df=45; Wolfskill vs Malathion_Resis2: adjusted p<0.0001, q=15.84, df=45). These lines were also deemed resistant when compared to the two new susceptible isofemale lines derived from the same population, Malathion_Sus1 and Malathion_Sus2 (Malathion_Sus1 vs Malathion_Resis1: adjusted p<0.0001, t=14.52, df=45; Malathion_Sus1 vs Malathion_Resis2: adjusted p<0.0001, t=16.78, df=45; Malathion_Sus2 vs Malathion_Resis1: adjusted p<0.0001, t=16.21, df=45; Malathion_Sus2 vs Malathion_Resis2: adjusted p<0.0001, t=18.48, df=45).

**Figure 2:**
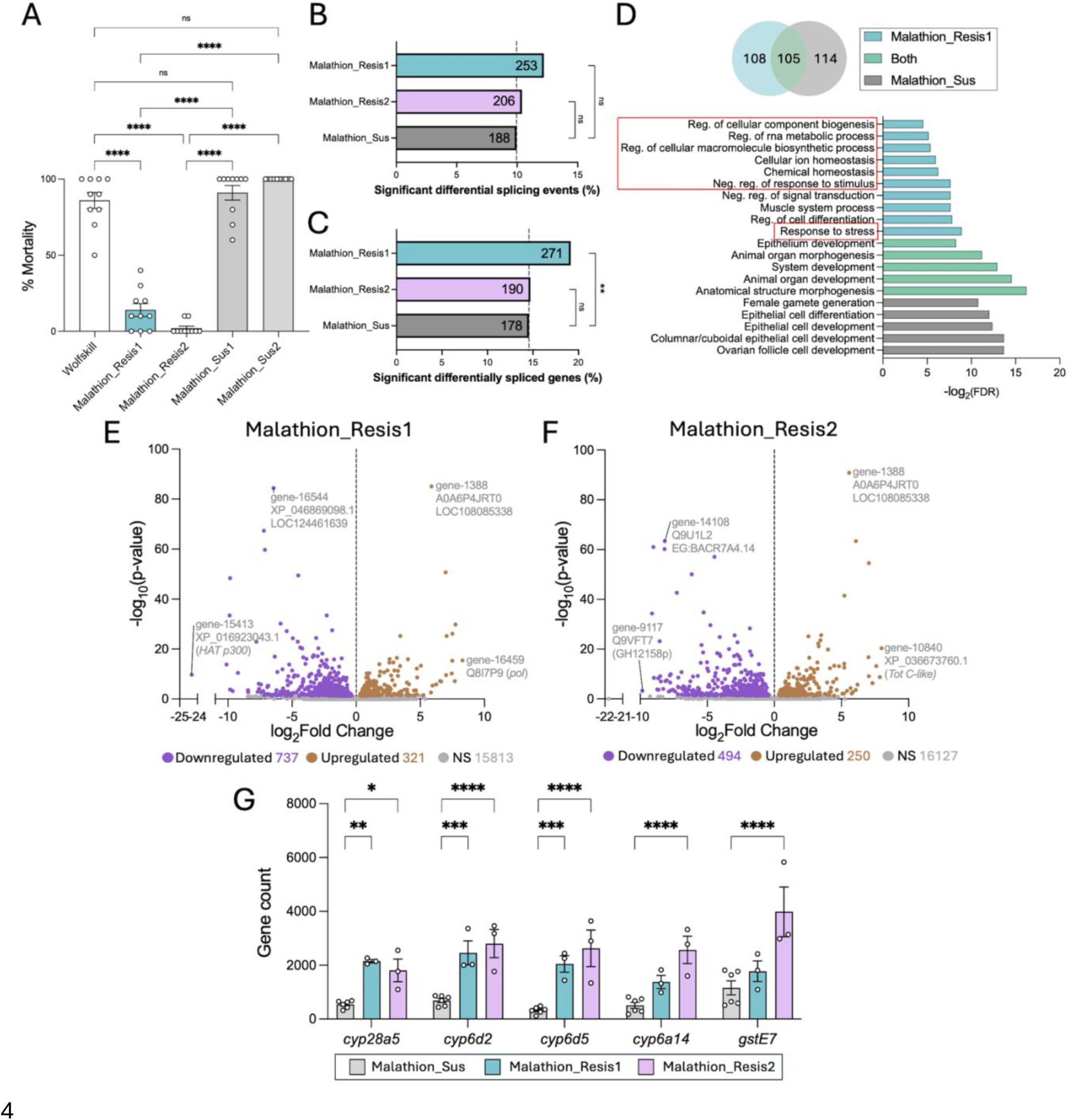
Increased alternative splicing is a common feature of metabolic resistance to Malathion, an additional insecticide. (A) Bioassay data at the discriminating dose (102.78ppm) revealing the resistance or susceptibility status of isofemale lines generated from Malathion-resistant populations collected in the field. An isofemale line from an untreated orchard served as the susceptible control (Wolfskill: white). Each point represents one replicate (n) that includes 5 males and 5 females (n=10), and error bars indicate ± SEM. The Malathion-resistant lines are in blue (Malathion_Resis1) and pink (Malathion_Resis2) while the susceptible lines are in grey. Asterisks denote significant p-values as determined by one-way ANOVA followed by Tukey’s multiple comparison test: **** p<0.0001. (B) Percentage of significant differential alternative splicing (AS) events out of all detected AS events in Malathion-resistant lines (Malathion_Resis1 (blue) or Malathion_Resis2 (pink)) vs Malathion-susceptible lines (Malathion_Sus1 and Malathion_Sus2). The number of significant AS events is displayed within each bar. The dotted line marks the % of significant differential splicing events in the susceptible group. Significance was determined by Fisher’s Exact test, corrected with Benjamini and Hochberg: **p<0.01. (C) Percentage of significant differentially spliced genes of all detected alternatively spliced genes in Malathion-resistant vs -susceptible lines. The number of significant AS genes is displayed within each bar. The dotted line marks the % of significant differentially spliced genes in the susceptible group. (D) Top: Venn diagram summarizing the number of processes enriched within the Gene Ontology (GO) Biological Processes categories for the differentially spliced genes in Malathion_Resis1 vs both Malathion-susceptible lines. Bottom: Categories that are the top 5 most significant in the resistant line alone (blue), the susceptible line alone (grey), and shared categories between the resistant and susceptible lines (green) or are categories that are related to stress response or processes previously implicated in resistance (red box), for instance, metabolic processes. The x-axis is the -log_2_(FDR), where FDR is the false discovery rate of enrichment p-values. Only the -log_2_(FDR) of the resistant line is shown for enriched categories in both the resistant and susceptible lines. (E) Gene expression differences between Malathion_Resis1 and the Malathion susceptible lines (Malathion_Sus1 and Malathion_Sus2 combined). Genes upregulated in the resistant population (brown) are to the right of the dotted line while downregulated genes (purple) lie to the left of the dotted line. Genes that exhibit no significant difference (NS) in expression between the resistant and susceptible lines are in grey. Labeled points signify genes that have either the largest fold change difference between the two groups or are the most significant. Labels contain the *D. suzukii* gene symbol (gene-XXXXX), the database ID, and the corresponding annotated gene symbol. (F) Gene expression differences between Malathion_Resis2 and the Malathion-susceptible lines. (G) Gene counts, or the number of reads per gene, of four *cytochrome P450* genes (*Cyp*) and *glutathione-s-transferase E7* (*GstE7*) in the Malathion-susceptible (grey) lines and the Malathion-resistant (Malathion_Resis1: blue; Malathion_Resis2: pink) isofemale lines. Each point denotes a biological replicate (n=3 replicates of 8-10 females each). Significance was determined by 2-way ANOVA followed by Sidak’s multiple comparisons test: * p<0.05, ** p<0.01, ***p<0.001, and **** p<0.0001.

We then subjected each resistant line (Malathion_Resis1 or Malathion_Resis2) to isoform sequencing and assessed splicing differences between the two resistant lines and the pooled susceptible lines (Malathion_Sus1 and Malathion_Sus2) (**Fig. 2B-C; S. Tables 16-18**). We noted that both resistant line comparisons did not have a significant difference in the number of differential splicing events as compared to the susceptible baseline comparison (Malathion_Sus1 versus Malathion_Sus2) (Malathion_Resis1 Fisher’s Exact Test=0.02603, q=0.05206; Malathion_Resis2 Fisher’s Exact Test=0.6767, q=0.6707). However, the Malathion_Resis1 comparison had more genes that were differentially spliced than the susceptible baseline comparison (Malathion_Resis1 Fisher’s Exact Test=0.001821, q=0.003642; Malathion_Resis2 Fisher’s Exact Test=0.9103, q=0.9103). We then performed functional enrichment analyses and observed that, similar to the Entrust- and Mustang Maxx-resistant lines (**Fig. 1E-H**), differentially spliced genes in the Malathion_Resis1 comparison are enriched in stress response pathways and pathways previously implicated in resistance, such as *regulation of cellular component biogenesis*, *chemical homeostasis*, and *response to stress* (**Fig. 2D; S. Tables 19-20**). No enriched processes for the differentially spliced genes were observed for the Malathion_Resis2 line.

We next sought to characterize the molecular mechanisms underlying Malathion resistance in these isofemale lines to determine if Malathion_Resis1, the line that exhibits aberrant splicing, also displays metabolic resistance. This would align with our findings so far that there is a strong association between aberrant splicing and metabolic resistance. We performed differentially expressed gene (DEG) analysis and observed a total of 1,058 DEGs in Malathion_Resis1, with 737 downregulated and 321 upregulated genes (**Fig. 2E; S. Table 21**). In Malathion_Resis2, there were 744 DEGs, with 494 downregulated and 250 upregulated (**Fig. 2F; S. Table 22**). Of the genes upregulated in the resistant lines, we observed metabolic genes previously implicated in insecticide resistance (**Fig. 2G**). For instance, both Malathion-resistant lines show increased expression of *cyp28a5* (Malathion_Resis1: p=0.0015, t=3.608, df=45; Malathion_Resis2: p=0.0137, t=2.834, df=45), *cyp6d2* (Malathion_Resis1: p=0.0005, t=3.971, df=45; Malathion_Resis2: p<0.0001, t=4.732, df=45), and *cyp6d5* (Malathion_Resis1: p=0.0007, t=3.853, df=45; Malathion_Resis2: p<0.0001, t=5.157, df=45), while only Malathion_Resis2 exhibited increased expression of *cyp6a14* (Malathion_Resis1: p=0.1112, t=1.952, df=45; Malathion_Resis2: p<0.0001, t=4.620, df=45) and *gstE7* (Malathion_Resis1: p=0.3224, t=1.372, df=45; Malathion_Resis2: p<0.0001, t=6.323, df=45). We did not observe known mutations that contribute to target-site resistance (Magaña et al. 2008; Perera et al. 2008). Taken together, the data suggest that Malathion resistance in these lines are attributed to metabolic resistance and hence changes in AS are associated with metabolic resistance in at least one additional resistant fly line. Thus, AS may underlie the development of insecticide resistance, or it could represent a response upon insecticide exposure.

### Alternative splicing is an adaptive response and may be a bet-hedging strategy

Organisms might use diverse strategies to adapt to environmental challenges, shaping populations as those strategies are selected across generations. In the context of insecticide resistance, alternative splicing could be used to diversify the genome, potentially increasing the chance for flies to survive insecticide exposure. This transcriptome-wide strategy is risky as some changes will be beneficial while others are not, incurring a high fitness cost, a preemptive measure known as bet-hedging (Murphy 1968; Nussbaum 1981). If a bet-hedging strategy is at play, the organisms that already carry an alternative splicing load, Malathion_Resis1 (**Fig. 2B-C**) in this case, would be equipped to respond to acute exposure to Malathion without any additional changes in AS. Conversely, those lines that do not bear this AS load, would react to insecticide exposure by actively changing their AS landscape. To test this hypothesis, we sought to expose the lines to sublethal doses of Malathion to evaluate the changes in the AS landscape (**Fig. 3**). First, we identified a sublethal insecticide dosage that both the resistant and susceptible flies could be exposed to without perishing (**Fig. 3A**). Using the Wolfskill susceptible line as a control, we tested four doses of Malathion (5ppm, 10ppm, 20ppm, and 40ppm) that were well below the resistance-discriminating dose of 102.78ppm (Disi et al. 2020). We identified 20ppm as a sublethal dosage of Malathion that susceptible flies would survive but might still be sufficient to elicit a physiological response.

**Figure 3:**
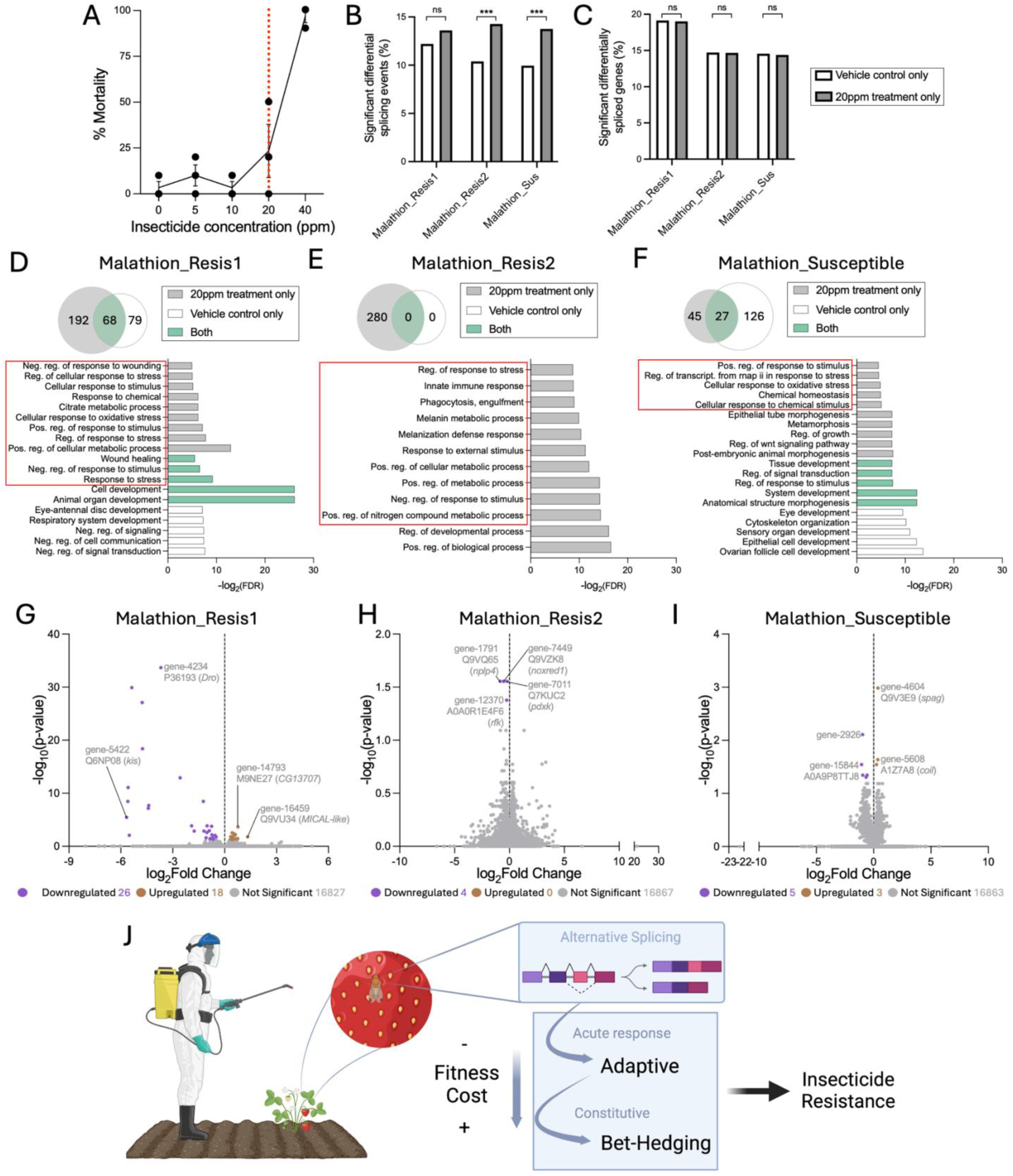
Aberrant splicing can be a response to Malathion treatment or be a bet-hedging strategy. (A) Mortality (%) of the Wolfskill population, a susceptible *D. suzukii* population, at various concentrations of Malathion. Each point represents a biological replicate (n=3 replicates of 5 females and 5 males each). The dotted red line denotes the sublethal concentration (20ppm) used for the remainder of the study. (B) Percentage of significant differential splicing events out of all AS events in the Malathion-resistant (Malathion_Resis1 or Malathion_Resis2) and Malathion-susceptible (Malathion_Sus1 and Malathion_Sus2) lines when treated with either 20ppm Malathion (grey) or the vehicle control (white). Significance was determined by Fisher’s Exact test, corrected with Benjamini and Hochberg: *** p<0.01. (C) Percentage of significant differentially spliced genes out of all detected AS genes in Malathion-resistant and -susceptible lines when treated with either 20ppm Malathion or the vehicle control. (D-F) Top: Venn diagram summarizing the number of processes enriched within the Gene Ontology (GO) Biological Processes categories for the differentially spliced genes in either (D) Malathion_Resis1, (E) Malathion_Resis2, or (F) the pooled Malathion-susceptible lines when treated with either a sublethal dosage of Malathion or the vehicle control. Bottom: Categories that are the top 5 most significant in the Malathion treatment alone (grey), the vehicle control alone (white), and shared categories between both conditions (green) or are categories that are related to stress response or processes previously implicated in resistance (red box), for instance, metabolic processes. The x-axis is the - log_2_(FDR), where FDR is the false discovery rate of enrichment p-values. Only the - log_2_(FDR) of the Malathion treatment is shown for enriched categories in both conditions. (G-I) Gene expression differences in (G) Malathion_Resis1, (H) Malathion_Resis2, and (I) the pooled Malathion susceptible lines (Malathion_Sus2 and Malathion_Sus1) when exposed to a sublethal dosage of Malathion vs the vehicle control. Genes upregulated in the groups treated with Malathion (brown) are to the right of the dotted line, while downregulated genes (purple) lie to the left of the dotted line. Genes that exhibit no significant difference (NS) in expression between the insecticide treatment and the vehicle control are in grey. Labeled points signify genes that have either the largest fold change difference between the insecticide treatment and the vehicle control or are the most significant, with the exception of (H) in which all DEG are labelled. Labels contain the *D. suzukii* gene symbol (gene-XXXXX), the database ID, and the corresponding annotated gene symbol. (J) A model proposing that global changes in AS occur in response to insecticide exposure to promote insecticide resistance. Because many genes are targeted, some of these changes are beneficial while others are not, resulting in fitness costs. Over time, AS is constitutively high, a bet-hedging, defensive strategy that flies use to protect themselves in the event of insecticide exposure.

We then surveyed the effect of sublethal Malathion exposure on the global AS patterns in the Malathion-resistant and -susceptible lines (**Fig. 3B-F; S. Tables 23-28**). We observed that Malathion_Resis2 and the Malathion-susceptible lines underwent more differential AS events when treated with insecticide while Malathion_Resis1 did not (Malathion_Resis1 Fisher’s Exact Test=0.1853, q=0.1853; Malathion_Resis2 Fisher’s Exact Test=1.463e-4, q=3.54e-4; Malathion Susceptible Fisher’s Exact Test=2.36e-4, q=3.54e-4) (**Fig. 3B; S. Tables 23-25**). Thus, Malathion_Resis1, which displays high levels of AS even in the absence of Malathion treatment (**Fig. 2C** and **3B**), appears to be preemptively responding to treatment whereas Malathion_Resis2, which does not show a high baseline of AS (**Fig. 2C** and **3B**), responds to Malathion treatment with an increase in AS events. This suggests that Malathion resistance is fixed in Malathion_Resis1, possibly due to bet-hedging, whereas Malathion resistance in Malathion_Resis2 may still be developing. Similarly, the pooled Malathion-susceptible lines also respond to acute Malathion treatment with an increase in AS events. In all 3 comparisons, Malathion treatment did not affect the total number of genes that were differentially spliced (Malathion_Resis1 Fisher’s Exact Test=0.9619, q=1.00; Malathion_Resis2 Fisher’s Exact Test=1.00, q=1.00; Malathion Susceptible Fisher’s Exact Test=0.9089, q=1.00) (**Fig. 3C; S. Tables 23-25**). However, the identity of the differentially spliced genes changed (**Fig. 3D-F; S. Tables 26-28**). We noted that the alternatively spliced genes in Malathion_Resis1 with and without sublethal Malathion exposure are enriched in stress response processes such as *wound healing* and *response to stress* (**Fig. 3D**). Following Malathion treatment, the AS genes were enriched in processes pertaining not only to stress response but also metabolism, such as *citrate metabolic process* and *positive regulation of cellular metabolic process*, which aligns with our finding that this line displays metabolic resistance (**Fig. 2E** and **G**). In Malathion_Resis2 exposed to the vehicle control, the differentially spliced genes were not enriched in any biological processes, however, following Malathion treatment, the AS genes were enriched in stress response processes, such as *regulation of response to stress* and *innate immune response*, and metabolic processes, such as *positive regulation of metabolic process* (**Fig. 3E**), consistent with the detection of metabolic resistance in Malathion_Resis2 (**Fig. 2F-G**). Interestingly, we also see an enrichment of cuticle-related processes such as *melanization defense response* and *melanin metabolic process* (**Fig. 3E**), suggesting that Malathion_Resis2 may not be fully committed to metabolic resistance. Rather, it may be resistant to Malathion via both penetration and metabolic resistance in the event of insecticide exposure. Finally, in the pooled Malathion-susceptible lines following Malathion treatment, the differentially spliced genes are enriched in stress response processes such as *cellular response to chemical stimulus* and *chemical homeostasis* (**Fig. 3F**), suggesting that AS is one way in which organisms adapt to environmental stress, more specifically, chemical exposure.

Furthermore, similar to what we observed in the Entrust and Mustang Maxx-resistant lines (**S. Fig 1**), the increase in AS did not appear to favor a particular type of AS (**S. Fig. 2**). Finally, the sublethal Malathion treatment had little to no effect on differential gene expression (**Fig. 3G-I; S. Tables 29-31**). Briefly, in Malathion_Resis1, Malathion treatment resulted in only 44 DEGs while treatment of Malathion_Resis2 resulted in only 4 DEGs, and in the susceptible lines, treatment resulted in only 8 DEGs. Taken together, our results indicate that sublethal Malathion treatment results in transcriptome-wide changes in AS, independent of resistance status, and remarkably, these changes occur without differential gene expression.

## Discussion

Alternative splicing (AS) results in various RNA transcripts that can produce different protein isoforms varying in structure and/or function. It is well-established that AS plays a role in adaptive evolution (Singh and Ahi 2022), and there is accumulating evidence showing that AS enables adaptations to extreme environmental conditions such as cold temperatures (reviewed in (Shiina and Shimizu 2020)) and chemical exposure (Zhu et al. 2017; Zhang et al. 2019; Wei et al. 2022). However, many studies investigated the AS of only a few key genes related to each adverse condition. Here, we explored whether changes in AS occur transcriptome-wide to potentially produce favorable adaptations, focusing on insecticide as the environmental stressor of interest and insecticide resistance and response as the adaptive traits.

Using a combination of short- and long-read sequencing, we assessed the AS landscape in isofemale lines of the keystone fruit pest, *Drosophila suzukii*, resistant to one of three classes of insecticides: spinosyns, pyrethroids, or organophosphates. We observed a general phenomenon in which lines showing metabolic resistance to insecticides exhibit an increase in the number of global AS events as well as the number of genes that are differentially spliced. Most importantly, challenging flies with a sublethal concentration of Malathion generated differential changes in the AS landscape, either in the number of genes, the number of events, or which processes the differentially spliced genes are enriched in. In particular, susceptible flies and resistant flies with comparable levels of AS to susceptible flies actively responded to sublethal insecticide exposure, plastically changing their AS landscapes, while another resistant line already exhibiting a high AS load in the absence of insecticide treatment did not. Thus, it is possible that AS is working as an immediate adaptive response to insecticide treatment such that many genes will be differentially spliced on the off chance that some of these changes will be beneficial for survival. In some cases, the affected genes will produce the resistant phenotype. Over time, these changes in AS will become heritable, so that even in the absence of insecticide treatment, the resistant individuals will exhibit high levels of AS to prepare for when they are exposed to insecticide, as a bet-hedging strategy (**Fig. 3J**). Taken together, our results suggest that AS may be an additional molecular mechanism underlying insecticide resistance.

Bet-hedging in insects has been linked to many adaptations to factors such as extreme climates (Rajon et al. 2014; Kain et al. 2015; Le Lann et al. 2021; van Baaren et al. 2024). Here, we show that this strategy could arise from a general stress response in the form of changes to the AS landscape. If true, this would prompt other interesting questions. For instance, are the patterns of the AS landscape (i.e., specific splicing events) inherited such that resistance becomes fixed at the population level? Or is the overactivation of AS a more general process that is being passed on through generations? Some possible mechanisms enabling the inheritance of AS across generations include epigenetic changes and mutations that affect splicing factor expression and/or recruitment (Pacini and Koziol 2018). Epigenetic changes have been linked to insecticide resistance, and there is accumulating evidence suggesting that histone modifications and chromatin organization can affect pre-mRNA splicing (Luco et al. 2011; Khan et al. 2012). There is also evidence suggesting that epigenetic changes are linked to insecticide resistance (Mogilicherla and Roy 2023). Thus, it is possible that AS is initially engaged as a stress response that gets refined over generations as epigenetic changes take place, and these changes then become passed on. This selection would result in stable resistant lines constitutively expressing a high AS load.

Interestingly, we noted that although there was an increase in AS in Malathion-treated flies, the sublethal concentration used was not sufficient to drive changes in gene expression (**Fig. 3G-I**). A previous study (Mishra et al. 2018) reported that insecticide treatment increases DEG in *D. suzukii*. This discrepancy is likely due to the lower concentration of insecticide (LC20), and the use of resistant flies, while Mishra et al. (2018) used the LC50 and susceptible flies. Regardless, the concentrations used here are of significant relevance for agriculture and biomedicine since recent surveys of water and soils have reported the presence of insecticide residues in the range of these concentrations, even after spraying has ceased (Del Prado-Lu 2015; Fisher et al. 2021; Kumari et al. 2022; Elliott et al. 2024; Larsen et al. 2024). Considering that the effects of the sublethal concentrations of insecticides detected here are at the splicing level, these otherwise undetected effects may have been previously overlooked when isoform sequencing was not performed. Our results add to the growing body of evidence showing the harmful effects of insecticide residues on the ecological landscape. Sublethal concentrations of insecticides and herbicides have been linked to the decline of the populations of other non-target species, including pollinators, such as wild bees, honey bees, and butterflies (Motta et al. 2018; Almeida et al. 2021; Siviter et al. 2021; Stuligross and Williams 2021; Gonzalez et al. 2022; Ward et al. 2022; Williams and Hemberger 2023; Gandara et al. 2024; Edwards et al. 2025).

The effects of insecticides extend not only to insects but also to other animals, including humans (Kaur et al. 2024). Several studies have linked insecticide exposure to cancer and long-term health complications (Navarrete-Meneses et al. 2017; Abolhassani et al. 2019; Ataei and Abdollahi 2022). The mechanisms behind these effects have been reported as widespread, from oxidative stress to changes in histone modifications (Kisby et al. 2009; Paul et al. 2018; Furlong et al. 2020). Given the association between AS and cancer (Bonnal et al. 2020; Sciarrillo et al. 2020; Zhang et al. 2021; Malhan et al. 2022; Kwok et al. 2025) as well as the association between cancer incidence and pesticide usage (Sun et al. 2020; Zhang et al. 2022), it is possible that exposure to insecticides, even at low dosages, as shown here, also promotes differential splicing in humans. Thus, this study provides insight into another molecular effect that insecticide exposure potentially has on humans.

Insecticide resistance is generally associated with high fitness costs (Kliot and Ghanim 2012). While we observed that the differentially spliced genes are enriched in processes associated with insecticide resistance, many other genes in biological processes necessary for proper cellular function and homeostasis are also affected in insecticide-resistant fly lines (**Fig. 1E-H**, **Fig. 2D**, and **Fig. 3D-F, S. Tables 10-15, 19-20,** and **26-28**). Thus, an unintended consequence of the AS landscape rearrangement to achieve resistance may be a decrease in fitness triggered by this non-specific response. This, and the observed AS response to acute insecticide treatment, suggests that global changes in AS may be a general stress reaction and not specific to just insecticide response and resistance. If that is the case, other stressors would also elicit high global changes in AS as an adaptive response. In the literature, it is well-supported that many abiotic factors, including temperature (reviewed in (Shiina and Shimizu 2020)) and nutrient stress (reviewed in (Ravi et al. 2015)), lead to changes in AS. However, whether AS is also being used in these contexts as a bet-hedging strategy and/or an acute adaptive stress response is unknown. Future studies can explore whether changes in the AS landscape are a general stress response, functioning as a common mechanism regardless of the stressor, or whether in these cases, it is directed to specific genes.

In addition to uncovering the association between global AS and insecticide resistance, this study could provide insights into the development of new classes of insecticides. Approaches to target alternative splicing have been taken to tackle human diseases, including new cancer treatments (Havens and Hastings 2016; Di et al. 2019). More so, since global changes in AS are associated with metabolic resistance, these newly designed classes of insecticides can presumably be used with many crops and a wider range of pests, decreasing the need for increased spraying of less specific and presumably more harmful chemicals.

Of important note, although we observe an association between resistance and global changes in AS, we are certainly not ruling out that there are other mechanisms involved that contribute to resistance development. For instance, regulation by miRNA (reviewed in (Mahalle et al. 2024)) or epigenetic regulation (reviewed in (Shilpa et al. 2024)) can play a role in resistance. Furthermore, we acknowledge that the changes we are observing are at the RNA level and recognize that high mRNA expression of an isoform does not necessarily mean that the protein isoform will also be highly expressed, given that there are many processes regulating the translation of mRNA to protein (reviewed in (Das et al. 2021; Brito Querido et al. 2024; Göransson and Strömblad 2024; Wu et al. 2024)). Further studies focusing on the extent of these differential AS events at the protein levels are warranted. Yet, in summary, this study advances our understanding of AS as a response to xenobiotic stress in both target and non-target species.

## Materials and Methods

### Field *Drosophila suzukii* populations and development of isofemale lines

The *Drosophila suzukii* isofemale lines (susceptible and resistant lines) established from either Mustang ® Maxx^TM^ 0.8 EC 9.15% zeta-cypermethrin (FMC Corporation, Philadelphia, PA, USA) or Entrust® SC 22.5% spinosad A & D (Corteva Agriscience, Indianapolis, IN, USA) resistant field populations used in this study were described in Tabuloc et al. 2024 (Tabuloc et al. 2024). The two Entrust-resistant lines are named as Entrust_Resis1 and Entrust_Resis2, and the susceptible lines are named as Entrust_Sus1 and Entrust_Sus2. The same convention was used to name isofemale lines that are resistant and susceptible to the other insecticides in this study.

To assess resistance to Malathion Insect Spray Concentrate 50% malathion (Spectrum Group, St. Louis, MO, USA), isofemale lines were established from *D. suzukii* collected in October 2020 from a raspberry field in Santa Cruz County, CA, USA and a strawberry field in Monterey County, CA, USA. Each population was confirmed to be resistant using bioassays (described in the “Discriminating dose bioassays and identifying a sublethal dosage of insecticide” section). Each isofemale line was then established from a single wild-caught, gravid, non-insecticide-treated female from the population. The siblings were crossed to one another for 8 generations to establish each isofemale line as described in Tabuloc et al. 2024 (Tabuloc et al. 2024). Bioassays were then performed on each isofemale line to identify resistance status. Two malathion-susceptible and malathion-resistant isofemale lines were used for subsequent analyses.

### Discriminating dose insecticide bioassays and identification of a sublethal dosage

Discriminating dose bioassays were performed using a glass vial residue bioassay. The interior of a 20ml glass scintillation vial (Fisher Scientific, Pittsburgh, PA, USA) was coated with insecticide solution at the pre-determined discriminating dose (Van Timmeren et al. 2018; Van Timmeren et al. 2019). Excess insecticide was removed from the vial, and the vials were left upright in the fume hood to dry overnight. Three- to five-day old flies (5 males and 5 females per replicate for a total of 10 replicates) from each isofemale line were transferred to a vial containing insecticide residue. The vials were maintained at room temperature for the duration of the experiment. Control vials consisted of residue of the insecticide vehicle.

Susceptibility of the isofemale lines resistant to either Entrust or Mustang Maxx was assessed in (Tabuloc et al. 2024). Additionally, susceptibility of the established isofemale lines to Malathion was assessed at the discriminating dose of 102.78 mg/liter (Disi and Sial 2021). Mortality was scored 6 hours after exposure to Malathion residue. An isofemale line generated from a population of *D. suzukii* from an untreated orchard in Solano County, CA, USA (referred to as “Wolfskill”) served as our susceptible control (Gress and Zalom 2019; Tabuloc et al. 2024). Differences in mortality between each isofemale line and the Wolfskill susceptible control were assessed by one-way ANOVA followed by Tukey’s multiple comparison test using GraphPad Prism Version 10.3.0 (GraphPad Software, La Jolla, CA, USA).

To identify a sublethal dosage of Malathion, the Wolfskill susceptible control was exposed to various concentrations of insecticide residue (0ppm, 5ppm, 10ppm, 20ppm, and 40ppm). The insecticide was diluted in acetone, and the vials were prepared as discussed above. Mortality was assessed 6 hours after insecticide exposure.

### RNA extraction, library preparation, and high-throughput sequencing

Female flies were entrained at 25°C in 12-hour light:12-hour dark cycles for two full days. On day 3, flies were collected on dry ice at ZT16, which is 16 hours after lights-on. This time point was selected because *D. suzukii* exhibits a low level of *cytochrome P450* expression at this time (Hamby et al. 2013), enabling any overexpression to be more readily observed. Fly bodies were separated from heads using frozen metal sieves (Newark Wire Cloth Company, Clifton, NJ, USA). Eight to ten female bodies were homogenized using a Kimble® pellet pestle® Cordless Motor (DWK Life Sciences, Rockwood, TN, USA) and Kimble™ Kontes™ Pellet Pestle™ (DWK Life Sciences) in 300 μL of TRI reagent (Sigma, St. Louis, MO). 60 μL of 100% chloroform (Sigma) was added to the homogenized tissue and incubated at room temperature for 10 minutes. The upper aqueous layer was recovered after spinning down and transferred to a new microcentrifuge tube. RNA was precipitated with an equal volume of 100% isopropanol at −20°C overnight. After spinning down, the RNA pellet was washed with 70% ethanol once and allowed to air dry. The pellet was then resuspended in 20 μL 1X Turbo DNA-free kit buffer (Thermo Fisher Scientific, Waltham, MA, USA) and treated with Turbo DNase per manufacturer’s instructions. RNA quality was assessed with the Agilent 2100 Bioanalyzer system (Agilent Technologies, Santa Clara, CA, USA) and the Qubit RNA IQ kit (Invitrogen, Waltham, MA, USA) on a Qubit 4 Fluorometer (Invitrogen). RNA purity was measured on a Nanodrop 1000 (Thermo Fisher Scientific), and RNA quantity was measured with the Qubit RNA HS (high sensitivity) assay kit (Invitrogen).

Illumina short-read sequencing libraries were prepared with 1 μg of high-quality RNA and the TruSeq Stranded mRNA Library Prep Kit (Illumina, San Diego, CA, USA) according to the manufacturer’s protocol. A total of twenty-four libraries were prepared: three biological replicates of two Malathion-resistant lines and two susceptible lines treated with either the acetone vehicle or 20ppm Malathion. Library insert size and quality were measured with the Agilent 2100 Bioanalyzer System, and library concentration was measured with the Qubit 4 Fluorometer. All libraries generated from *D. suzukii* lines treated with the acetone vehicle were pooled together, and all libraries generated from *D. suzukii* lines treated with the sublethal dosage of Malathion were pooled together, such that there were twelve libraries per pool. Pooled samples were sent to Novogene (Sacramento, CA, USA) for sequencing on the HiSeq X Ten platform (Illumina) using PE150.

For PacBio Iso-Seq, high-quality RNA was sent to the DNA Technologies and Expression Analysis Core Laboratory at UC Davis for library preparation and sequencing. A total of 48 libraries were sequenced on five 8M SMRT cells on a PacBio Sequel system (PacBio, Menlo Park, CA, USA). Each of the five SMRT cells consisted of one of the following pooled libraries: (1) all Entrust resistant and susceptible lines, (2) all Mustang Maxx resistant and susceptible lines, and (3-5) 1 of 3 biological replicates of vehicle- and Malathion-treated *D. suzukii* that are either resistant or susceptible to Malathion.

### Repeat element identification and genome re-annotation

We used the NCBI *Drosophila suzukii* Annotation Release 102 based on the LBDM_Dsuz_2.1.pri assembly (accession no. GCF_013340165.1) (Paris et al. 2020) to generate a *de novo* repeat library using RepeatModeler2 (v 2.0.4) (Flynn et al. 2020) using default settings and ‘-LTRStruct’ to predict long terminal repeats (LTRs). We then softmasked the genome assembly using the Dfam v 3.7 database with RepeatMasker (v 4.1.5) (Chen 2004), followed by a second round of softmasking using RepeatMasker (v 4.1.5) and the *de novo* repeat library we previously generated. We identified and masked 135,904,552 bp of repeat regions in the reference genome, accounting for 50.7% of the assembly (**S. Table 1**), in close agreement with a previous report (Mérel et al. 2021). Among classes of repeat elements, short interspersed nuclear elements (SINE) accounted for 20.21% of all repeats.

Next, we mined NCBI’s Sequence Read Archive (SRA) for publicly available *D. suzukii* RNAseq datasets (last accessed June 28th, 2023), compiling a list of 56 sequencing datasets spanning both sexes and multiple tissue types (**S. Table 2**). Downloaded SRAs and in-house short-read RNAseq libraries were first quality checked using fastqc (v 0.12.1, http://www.bioinformatics.babraham.ac.uk/projects/fastqc) and multiqc (v 1.14) (Ewels et al. 2016). Sequencing adapters and low-quality reads were trimmed using Trimmomatic (v 0.39) (Bolger et al. 2014) and then each library was quality-checked again using fastqc and multiqc. Cleaned short-reads were aligned to the assembly using Hisat2 (v 2.2.1) (Kim et al. 2019) and used as evidence along with the OrthoDB arthropod protein dataset (arthropoda_odb11, last accessed June 28th, 2023) and the *D. melanogaster* proteome (Uniprot proteome accession number UP000000803_7227) to train ProtHint, Genemark-ETP+, and AUGUSTUS implemented in BRAKER3 (v 3.0.3) (Gotoh 2008; Stanke et al. 2008; Iwata and Gotoh 2012; Buchfink et al. 2015; Hoff et al. 2016; Hoff et al. 2019; Kovaka et al. 2019; Pertea and Pertea 2020; Brůna et al. 2021; Bruna et al. 2024; Gabriel et al. 2024).

We then incorporated PacBio Iso-Seq long-reads into our gene model predictions by following the BRAKER3 ‘long_read_protocol’ (https://github.com/Gaius-Augustus/BRAKER/blob/master/docs/long_reads/long_read_protocol.md). Briefly, we pooled all demultiplexed Iso-Seq CCS reads together and ran the isoseq (v 4.0.0, Pacific Biosciences, USA) pipeline to generate full-length non-chimeric (FLNC) isoform sequences. FLNC sequences were then clustered using ‘isoseq cluster’ and the resulting high-quality transcript sequences were mapped to the softmasked assembly using pbmm2 (v 1.11.99) with default settings. Redundant mapped transcripts were collapsed using ‘isoseq collapse’ with default settings and ‘—do-not-collapse-extra-5exons’. Protein-coding regions of mapped Iso-Seq transcripts were predicted using GeneMarkS-T (v. 5.1) (Tang et al. 2015) and the results were converted to GTF format. Finally, the long-read version of TSEBRA (https://github.com/Gaius-Augustus/TSEBRA/tree/long_reads) was used to merge predicted gene models from the BRAKER3 run and Iso-Seq transcripts to generate a GTF file. AGAT (v. 1.2.0) (https://www.doi.org/10.5281/zenodo.3552717) was used to convert the GTF file to GFF format, collapse any remaining redundant transcripts, and annotate pseudogenes. Ribosomal RNAs and transfer RNAs were annotated using rnammer (v. 1.2) (Lagesen et al. 2007) and tRNAscan-SE (v. 2.0.12) (Chan et al. 2021), respectively. In total, we predicted 17,159 gene models including 954 protein-coding gene models not previously identified in the reference transcriptome. Our re-annotation resulted in 34,354 transcript models, increasing the number of previously predicted transcripts by 8,814 (**S. Table 3**) and the mean transcripts per gene from 1.6 to 2.1.

Functional annotations were then assigned to the final transcript set using EnTAP (v 1.0.1) (Hart et al. 2020) in ‘-runP’ mode using the RefSeq Invertebrate protein database, UniProt SwissProt database, UniProtKB/TrEMBL database, and the *D. melanogaster* proteome (Uniprot proteome accession number UP000000803_7227) (all last accessed February 14th, 2024). We assigned functional annotations to 92.41% of all transcript models.

### Alternative splicing analysis

Using Salmon (v. 1.10.1), cleaned Illumina RNA-Seq reads were mapped to the *Drosophila suzukii* genome assembly (RefSeq: GCF_013340165.1) and gene expression was quantified using default parameters (Patro et al. 2017). Differential expression of gene isoforms and splicing events were determined using SUPPA2 (v.2.3) (Trincado et al. 2018). Specifically, splicing events on the updated annotation were determined with generateEvents. Using the option ‘-e {SE,SS,MX,RI,FL}’, we opted to call exon skipping (SE), alternative splice sites (SS), mutually exclusive exons (MX), intron retention (RI), and first/last exons (FL). Relative abundance of each splicing event (psi) was determined with psiPerEvent. Finally, differential isoform expression was assessed with diffsplice using classical mode, using the pooled Susceptible lines as reference for Resistant vs. Susceptible and Sus_1 as reference for Susceptible vs. Susceptible comparisons. Unless otherwise stated, default parameters were used for all analyses done with SUPPA2.

Features with FDR < 0.05 were considered differentially expressed. The number of significant differentially spliced events for each group (either Resis1 versus pooled susceptible lines, Resis2 versus pooled susceptible lines, or Sus_1 versus Sus_2) was divided by all detected events and multiplied by 100 to get the percentage of significant differentially spliced events that occurred. Similarly, the number of genes that have at least one significant splicing event was divided by all detected genes and multiplied by 100 to get the percentage of significant differentially spliced genes that are present. The values between each resistant line and the pooled susceptible group for each insecticide were compared using Fisher’s Exact Test followed by Benjamini and Hochberg’s multiple comparison test on R. Both the values of the Fisher’s Exact Test and the corrected q value are provided throughout the manuscript.

### Functional enrichment analysis

Gene Ontology (GO) enrichment of genes was performed using ShinyGO 0.76.3 (Ge et al. 2020). GO terms and processes were considered enriched if the false discovery rate (FDR) < 0.05. GO categories for the enrichment performed on differential splicing data were divided into categories only enriched in the resistant line comparisons (either Resis1 versus pooled susceptible lines or Resis2 versus pooled susceptible lines), only in the susceptible line comparison (Sus1 versus Sus_2), or overlap between the two comparisons.

### Differential expression gene (DEG) analysis

Differential expression gene analysis for the Malathion-resistant and -susceptible lines was performed using sequencing reads derived from Illumina short-read sequencing. First, rRNA reads were removed using SortMeRNA (v. 2.1) (Kopylova et al. 2012). Adapters (ILLUMINACLIP parameters 2:30:10) and low-quality ends (LEADING: 10, TRAILING:10, MINLEN:36) were trimmed using fastp (v. 0.24.0) (Chen et al. 2018). Cleaned reads were aligned to our *Drosophila suzukii* genome reannotation based on the LBDM_Dsuz_2.1.pri assembly (accession no. GCF_013340165.1) (Paris et al. 2020) using STAR (v. 2.7.11b) (Dobin et al. 2013). Count data from STAR (--quantMode GeneCounts) served as input in the DESeq2 package (Love et al. 2014) in R to perform differential expression analysis on each resistant line vs both pooled susceptible lines. Each Malathion-resistant line was compared to pooled susceptible lines separately as each resistant line might exhibit resistance due to different mechanisms (Tabuloc et al. 2024).

Additional comparisons were performed to assess whether exposure to a sublethal dosage elicits a response at the gene expression level. Thus, the acetone vehicle was compared to the Malathion treatment. Both susceptible lines were pooled together in these comparisons since they were combined in the resistant to susceptible comparisons. For each comparison, genes with fold change differences between either resistant vs susceptible populations of vehicle vs insecticide treatment with a Benjamini-Hochberg adjusted p-value < 0.05 were considered differentially expressed. Correlation between biological replicates was calculated with Pearson’s correlation coefficient (**S. Fig. 3**), which was determined with the ‘stats’ package in R (v. 4.2.1). Expression differences of key genes between the resistant and susceptible lines were calculated with two-way ANOVA followed by two-stage linear set-up procedure of Benjamini, Krieger, and Yekutieli on GraphPad Prism.

## Acknowledgements

We thank all members of the Chiu laboratory and members of the SCRI spotted wing *Drosophila* research team, in particular Rufus Isaacs, Steven Van Timmeren, Elizabeth Beers, Gregory Loeb, Ashfaq Sial, and Vaughn Walton for their feedback and suggestions. We would like to give a special thank you to Dr. Yao Cai, whose feedback greatly improved the manuscript. This project is supported by the National Institute of Food and Agriculture at the United States Department of Agriculture, awards 2020-51181-32140 and CA-D-ENM-2150-H to JCC, and the CDFA Specialty Crop Block Grant 21SCBPCA1002 awarded to JCC.

## Authors Contributions

CAT and JCC conceived the study; JCC and FGZ supervised the study; CAT performed the experiments; CAT, SH, CRC, and JCC performed bioinformatic analyses with the input of HZ; CAT, SH, and JCC contributed to critical interpretation of the data; CAT and SH wrote the paper with the input of JCC and all other authors.

## Data availability and sharing plan

Raw Illumina and PacBio reads have been deposited to the NCBI Sequence Read Archive (SRA) and can be found under BioProject accession numbers PRJNA983428 and PRJNA1143070. Supplementary Tables 4-31 contain the lists of differentially spliced genes, differentially expressed genes, and the results of the functional enrichment analyses and have been uploaded to Dryad (https://datadryad.org/share/7xlDoiG7WR_2ClNUMbSWDa9jG2zP-Ky5_Ipsu2R6hGQ).

