## Supplemental Figs. 1-5 and Supplemental Tables 1 and 31 for "Global changes in alternative splicing are associated with insecticide response and resistance"

### Supporting information:

**Supplemental Figs. 1-5 and Supplemental Tables 1 and 3.**

**Note that the following are provided as .csv or .xlsx files on Dryad**
(https://datadryad.org/share/7xlDoiG7WR_2ClNUMbSWDa9jG2zP-Ky5_Ipsu2R6hGQ)

***S. Table 2:*** *List of sequencing datasets used to re-annotate the D. suzukii genome assembly*
***S. Table 4***: *List of alternative splicing (AS) events for Entrust_Resis1 vs. Entrust-susceptible lines*
***S. Table 5***: *List of alternative splicing (AS) events for Entrust_Resis2 vs. Entrust-susceptible lines*
***S. Table 6:*** *List of alternative splicing (AS) events for Entrust_Sus1 vs. Entrust_Sus2*
***S. Table 7***: *List of alternative splicing (AS) events for MustangMaxx_Resis1 vs. Mustang Maxx-susceptible lines*
***S. Table 8***: *List of alternative splicing (AS) events for MustangMaxx_Resis2 vs. Mustang Maxx-susceptible lines*
***S. Table 9:*** *List of alternative splicing (AS) events for MustangMaxx-Sus1 vs. MustangMaxx_Sus2*
***S. Table 10***: *Enrichment for alternative splicing (AS) events for Entrust_Resis1 vs. Entrust-susceptible lines*
***S. Table 11:*** *Enrichment for alternative splicing (AS) events for Entrust_Sus1 vs. Entrust_Sus2*
***S. Table 12***: *Enrichment for alternative splicing (AS) events for Entrust_Resis2 vs. Entrust-susceptible lines*
***S. Table 13***: *Enrichment for alternative splicing (AS) events for MustangMaxx_Resis1 vs. Mustang Maxx-susceptible lines*
***S. Table 14***: *Enrichment for alternative splicing (AS) events for MustangMaxx-Sus1 vs. MustangMaxx_Sus2*
***S. Table 15***: *Enrichment for alternative splicing (AS) events for MustangMaxx_Resis2 vs. Mustang Maxx-susceptible lines*
***S. Table 16***: *List of alternative splicing (AS) events for Malathion_Resis1 vs. Malathion-susceptible lines*
***S. Table 17***: *List of alternative splicing (AS) events for Malathion_Resis2 vs. Malathion-susceptible lines*
***S. Table 18:*** *List of alternative splicing (AS) events for Malathion_Sus1 vs. Malathion_Sus2*
***S. Table 19***: *Enrichment for alternative splicing (AS) events in Malathion_Resis1 vs. Malathion-susceptible lines*
***S. Table 20:*** *Enrichment for alternative splicing (AS) events in Malathion_Sus1 vs. Malathion_Sus2*
***S. Table 21***: *List of differentially expressed genes (DEGs) for Malathion_Resis1 vs. Malathion-susceptible lines*
***S. Table 22***: *List of differentially expressed genes (DEGs) for Malathion_Resis2 vs. Malathion-susceptible lines*

***S. Table 23:*** *List of alternative splicing (AS) events for Malathion_Resis1 vs. Malathion-susceptible lines when treated with 20ppm Malathion*
***S. Table 24:*** *List of alternative splicing (AS) events for Malathion_Resis2 vs. Malathion-susceptible lines when treated with 20ppm Malathion*
***S. Table 25:*** *List of alternative splicing (AS) events for Malathion_Sus1 vs. Malathion_Sus2 when treated with 20ppm Malathion*
***S. Table 26:*** *Enrichment for alternative splicing (AS) events for Malathion_Resis1 vs. Malathion-susceptible lines when treated with 20ppm Malathion*
***S. Table 27:*** *Enrichment for alternative splicing (AS) events for Malathion_Sus1 vs. Malathion_Sus2 when treated with 20ppm Malathion*
***S. Table 28:*** *Enrichment for alternative splicing (AS) events for Malathion_Resis2 vs. Malathion-susceptible lines when treated with 20ppm Malathion*
***S. Table 29:*** *List of differentially expressed genes (DEGs) in Malathion_Resis1 20ppm Malathion vs. vehicle control*
***S. Table 30:*** *List of differentially expressed genes (DEGs) in Malathion_Resis2 20ppm Malathion vs. vehicle control*
***S. Table 31:*** *List of differentially expressed genes (DEGs) in Malathion-susceptible lines 20ppm Malathion vs. vehicle control*

#### S. Table 1: Repeatome statistics for *Drosophila suzukii* transcriptome re-annotation

| Class | Number | Length (bp) | % of assembly |
|---|---|---|---|
| SINE | 463 | 138677 | 0.05 |
| LINE | 49006 | 23517681 | 8.77 |
| LTR | 102183 | 54168690 | 20.21 |
| DNA | 63863 | 19204013 | 7.16 |
| RC | 84496 | 14402743 | 5.37 |
| Other | 209856 | 9027445 | 3.38 |
| Unclassified | 54218 | 15444975 | 5.76 |
| Artifact | 5 | 328 | 0 |
| Total | 564090 | 135904552 | 50.7 |

SINE: short interspersed nuclear elements; LINE: long interspersed nuclear elements; LTR: long terminal repeats; DNA: DNA elements; RC: rolling circle elements; Other: small RNA, satellite DNA, simple repeats, and low complexity repeats; bp: base pair

**S. Table 3: Statistics for the updated *Drosophila suzukii* transcriptome**

| | **Updated annotation** | **Paris et. al 2020** |
|---|---|---|
| Number of genes | 17159 | 17539 |
| Number of transcripts | 34354 | 25540 |
| Number of protein-coding genes | 16521 | 15567 |
| Mean transcripts per gene | 2.1 | 1.6 |
| Mean exons per transcript | 4.7 | 6.5 |
| Mean gene length (bp) | 6719 | 7506 |
| Mean transcript length (bp) | 7770 | 14301 |
| Mean CDS length (bp) | 1782 | 2124 |
| Transcripts > 500bp | 28637 | 22223 |
| Transcripts > 1000bp | 21140 | 16708 |
| Longest transcript (bp) | 62712 | 53532 |
| Number of rRNA | 397 | 284 |
| Number of tRNA | 350 | 360 |

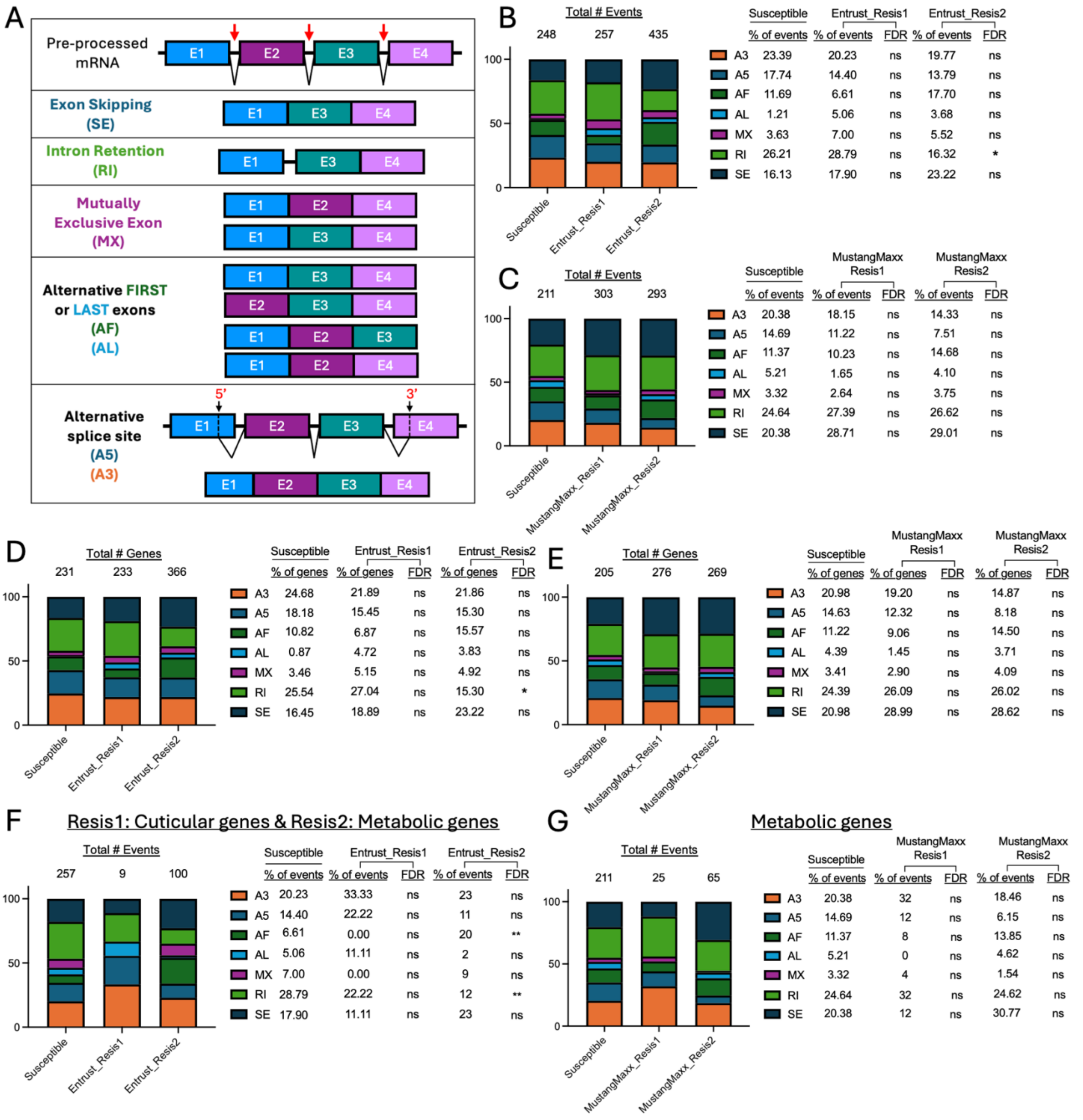


**S. Fig 1: Insecticide-resistant lines do not exhibit a specific type of alternative splicing.** (A) Schematic describing 5 different types of alternative splicing (AS) events, where the colored boxes represent exons and the black line represents introns. (B-C) Distribution of AS events in (B) Entrust-resistant (Entrust_Resis1 or Entrust_Resis2) and Entrust-susceptible lines (Entrust_Sus1 vs Entrust_Sus2) and (C)

Mustang Maxx-resistant (MustangMaxx_Resis1 and MustangMaxx_Resis2) and Mustang Maxx-susceptible lines, categorized by AS type described in panel A. The total number of significant AS events for each line is noted at the top of each bar. The % of significant events for each AS type and the False Discovery Rate (FDR) comparing each resistant line to the susceptible control is noted to the right of the graph. Asterisks denote significance as determined by Fisher's Exact test, corrected with Benjamini-Hochberg: *$p<0.05$ and **$p<0.01$. (D-E) Distribution of genes exhibiting each AS event in (D) Entrust-resistant and Entrust-susceptible lines and (E) Mustang Maxx-resistant and Mustang Maxx-susceptible lines. The number of significant differentially spliced genes for each line is noted at the top of each bar. The % of significant differentially spliced genes for each AS type and the FDR comparing each resistant line to the susceptible control is noted to the right of the graph. (F) Distribution of AS events in genes involved in either cuticle-related or metabolic processes in Entrust-resistant lines and AS events of all genes in Entrust-susceptible lines. (G) Distribution of AS events in genes involved in metabolic processes in Mustang Maxx-resistant lines and AS events of all genes in Mustang Maxx-susceptible lines.

**A**

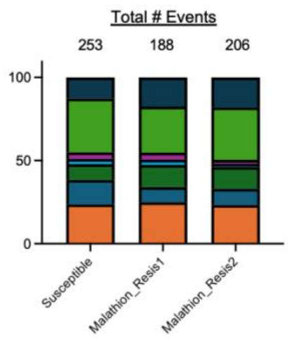


| | Vehicle | | | | |
|---|---|---|---|---|---|
| | Susceptible | Malathion Resis1 | | Malathion Resis2 | |
| | % of events | % of events | FDR | % of events | FDR |
| A3 | 23.72 | 25.00 | ns | 23.30 | ns |
| A5 | 14.62 | 9.04 | ns | 9.71 | ns |
| AF | 9.49 | 13.30 | ns | 13.11 | ns |
| AL | 3.16 | 3.19 | ns | 1.94 | ns |
| MX | 3.95 | 4.26 | ns | 2.43 | ns |
| RI | 32.41 | 27.66 | ns | 31.55 | ns |
| SE | 12.65 | 17.55 | ns | 17.96 | ns |

**B**

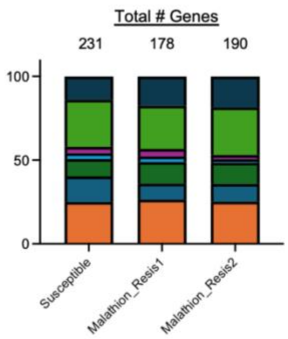


| | Vehicle | | | | |
|---|---|---|---|---|---|
| | Susceptible | Malathion Resis1 | | Malathion Resis2 | |
| | % of genes | % of genes | FDR | % of genes | FDR |
| A3 | 25.11 | 26.40 | ns | 25.26 | ns |
| A5 | 15.15 | 9.55 | ns | 10.53 | ns |
| AF | 10.39 | 12.92 | ns | 12.63 | ns |
| AL | 3.46 | 3.37 | ns | 2.11 | ns |
| MX | 3.90 | 4.49 | ns | 2.63 | ns |
| RI | 28.14 | 25.84 | ns | 28.42 | ns |
| SE | 13.85 | 17.42 | ns | 18.42 | ns |

**C**

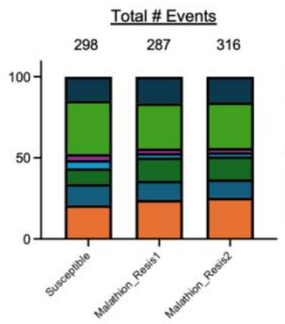


| | Insecticide | | | | |
|---|---|---|---|---|---|
| | Susceptible | Malathion Resis1 | | Malathion Resis2 | |
| | % of events | % of events | FDR | % of events | FDR |
| A3 | 20.81 | 25.00 | ns | 23.30 | ns |
| A5 | 13.09 | 9.04 | ns | 9.71 | ns |
| AF | 9.73 | 13.30 | ns | 13.11 | ns |
| AL | 5.03 | 3.19 | ns | 1.94 | ns |
| MX | 3.69 | 4.26 | ns | 2.43 | ns |
| RI | 32.89 | 27.66 | ns | 31.55 | ns |
| SE | 14.77 | 17.55 | ns | 17.96 | ns |

**D**

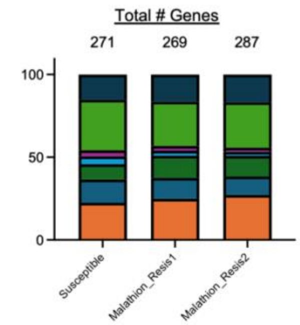


| | Insecticide | | | | |
|---|---|---|---|---|---|
| | Susceptible | Malathion Resis1 | | Malathion Resis2 | |
| | % of genes | % of genes | FDR | % of genes | FDR |
| A3 | 22.51 | 24.91 | ns | 27.18 | ns |
| A5 | 14.02 | 12.64 | ns | 11.15 | ns |
| AF | 9.23 | 13.82 | ns | 12.54 | ns |
| AL | 4.80 | 2.97 | ns | 2.44 | ns |
| MX | 3.69 | 2.97 | ns | 2.44 | ns |
| RI | 30.63 | 26.77 | ns | 27.53 | ns |
| SE | 15.13 | 16.36 | ns | 16.72 | ns |

**S. Fig 2: Exposure to a sublethal dosage of insecticide does not result in an enrichment in a particular alternative splicing type.** (A-B) Distribution of (A) AS events and (B) genes exhibiting each AS type (described in **S. Fig. 1A**) in Malathion-resistant (Malathion_Resis1 and Malathion_Resis2) and Malathion-susceptible lines treated with the vehicle control. The number of significant (A) AS events and (B) differentially spliced genes for each line is noted at the top of the bars. The % of significant (A) events and (B) genes for each AS type and the False Discovery Rate (FDR) comparing each resistant line to the susceptible control is noted to the right of the graph. Significance was determined by Fisher's Exact test, corrected with Benjamini-Hochberg. (C-D) Distribution of (C) AS events and (D) genes exhibiting each AS type in Malathion-resistant and Malathion-susceptible lines treated with a sublethal dosage of Malathion.

| | | | Resistant | | | | | | | | | | | | Susceptible | | | | | | | | | | | |
|---|---|---|---|---|---|---|---|---|---|---|---|---|---|---|---|---|---|---|---|---|---|---|---|---|---|---|
| | | | Vehicle | | | Insecticide | | | Vehicle | | | Insecticide | | | Vehicle | | | Insecticide | | | Vehicle | | | Insecticide | | |
| | | | Resis1_rep1 | Resis1_rep2 | Resis1_rep3 | Resis1_rep1 | Resis1_rep2 | Resis1_rep3 | Resis2_rep1 | Resis2_rep2 | Resis2_rep3 | Resis2_rep1 | Resis2_rep2 | Resis2_rep3 | Sus1_rep1 | Sus1_rep2 | Sus1_rep3 | Sus1_rep1 | Sus1_rep2 | Sus1_rep3 | Sus2_rep1 | Sus2_rep2 | Sus2_rep3 | Sus2_rep1 | Sus2_rep2 | Sus2_rep3 |
| Resistant | Vehicle | Resis1_rep1 | 1.0000 | | | | | | | | | | | | | | | | | | | | | | | |
| | | Resis1_rep2 | 0.9850 | 1.0000 | | | | | | | | | | | | | | | | | | | | | | |
| | | Resis1_rep3 | 0.9673 | 0.9723 | 1.0000 | | | | | | | | | | | | | | | | | | | | | |
| | Insecticide | Resis1_rep1 | 0.9894 | 0.9894 | 0.9640 | 1.0000 | | | | | | | | | | | | | | | | | | | | |
| | | Resis1_rep2 | 0.9867 | 0.9909 | 0.9708 | 0.9907 | 1.0000 | | | | | | | | | | | | | | | | | | | |
| | | Resis1_rep3 | 0.9941 | 0.9883 | 0.9667 | 0.9951 | 0.9938 | 1.0000 | | | | | | | | | | | | | | | | | | |
| | Vehicle | Resis2_rep1 | 0.9259 | 0.9223 | 0.9725 | 0.9172 | 0.9356 | 0.9263 | 1.0000 | | | | | | | | | | | | | | | | | |
| | | Resis2_rep2 | 0.9712 | 0.9719 | 0.9722 | 0.9653 | 0.9805 | 0.9695 | 0.9574 | 1.0000 | | | | | | | | | | | | | | | | |
| | | Resis2_rep3 | 0.9751 | 0.9704 | 0.9828 | 0.9735 | 0.9834 | 0.9801 | 0.9768 | 0.9806 | 1.0000 | | | | | | | | | | | | | | | |
| | Insecticide | Resis2_rep1 | 0.9812 | 0.9763 | 0.9661 | 0.9787 | 0.9894 | 0.9858 | 0.9502 | 0.9842 | 0.9892 | 1.0000 | | | | | | | | | | | | | | |
| | | Resis2_rep2 | 0.9651 | 0.9554 | 0.9669 | 0.9576 | 0.9605 | 0.9673 | 0.9451 | 0.9555 | 0.9787 | 0.9740 | 1.0000 | | | | | | | | | | | | | |
| | | Resis2_rep3 | 0.9575 | 0.9582 | 0.9773 | 0.9588 | 0.9654 | 0.9620 | 0.9798 | 0.9722 | 0.9855 | 0.9733 | 0.9587 | 1.0000 | | | | | | | | | | | | |
| Susceptible | Vehicle | Sus1_rep1 | 0.9606 | 0.9438 | 0.9644 | 0.9474 | 0.9656 | 0.9586 | 0.9652 | 0.9752 | 0.9808 | 0.9773 | 0.9601 | 0.9606 | 1.0000 | | | | | | | | | | | |
| | | Sus1_rep2 | 0.9467 | 0.9413 | 0.9609 | 0.9436 | 0.9613 | 0.9497 | 0.9697 | 0.9704 | 0.9745 | 0.9654 | 0.9279 | 0.9724 | 0.9822 | 1.0000 | | | | | | | | | | |
| | | Sus1_rep3 | 0.9783 | 0.9593 | 0.9507 | 0.9696 | 0.9780 | 0.9784 | 0.9401 | 0.9687 | 0.9780 | 0.9848 | 0.9534 | 0.9642 | 0.9812 | 0.9752 | 1.0000 | | | | | | | | | |
| | Insecticide | Sus1_rep1 | 0.9704 | 0.9504 | 0.9330 | 0.9616 | 0.9685 | 0.9743 | 0.9094 | 0.9459 | 0.9658 | 0.9765 | 0.9655 | 0.9283 | 0.9708 | 0.9405 | 0.9792 | 1.0000 | | | | | | | | |
| | | Sus1_rep2 | 0.9792 | 0.9787 | 0.9756 | 0.9783 | 0.9841 | 0.9832 | 0.9488 | 0.9794 | 0.9831 | 0.9844 | 0.9705 | 0.9722 | 0.9796 | 0.9739 | 0.9819 | 0.9739 | 1.0000 | | | | | | | |
| | | Sus1_rep3 | 0.9598 | 0.9386 | 0.9184 | 0.9465 | 0.9513 | 0.9620 | 0.9000 | 0.9269 | 0.9519 | 0.9632 | 0.9565 | 0.9163 | 0.9568 | 0.9247 | 0.9667 | 0.9917 | 0.9593 | 1.0000 | | | | | | |
| | Vehicle | Sus2_rep1 | 0.7795 | 0.8088 | 0.8392 | 0.7845 | 0.8287 | 0.7847 | 0.8662 | 0.8802 | 0.8366 | 0.8262 | 0.7576 | 0.8545 | 0.8443 | 0.8855 | 0.8099 | 0.7460 | 0.8244 | 0.7185 | 1.0000 | | | | | |
| | | Sus2_rep2 | 0.8501 | 0.8583 | 0.8937 | 0.8530 | 0.8544 | 0.8405 | 0.8881 | 0.9047 | 0.8643 | 0.8506 | 0.8050 | 0.8995 | 0.8658 | 0.9096 | 0.8487 | 0.7748 | 0.8721 | 0.7546 | 0.9083 | 1.0000 | | | | |
| | | Sus2_rep3 | 0.9117 | 0.9232 | 0.9587 | 0.9207 | 0.9205 | 0.9140 | 0.9470 | 0.9353 | 0.9392 | 0.9162 | 0.8991 | 0.9615 | 0.9200 | 0.9535 | 0.9115 | 0.8626 | 0.9376 | 0.8460 | 0.8782 | 0.9567 | 1.0000 | | | |
| | Insecticide | Sus2_rep1 | 0.8643 | 0.8852 | 0.9168 | 0.8646 | 0.9027 | 0.8691 | 0.9374 | 0.9408 | 0.9161 | 0.9050 | 0.8572 | 0.9253 | 0.9231 | 0.9466 | 0.8891 | 0.8448 | 0.9063 | 0.8250 | 0.9753 | 0.9234 | 0.9299 | 1.0000 | | |
| | | Sus2_rep2 | 0.9631 | 0.9532 | 0.9714 | 0.9574 | 0.9558 | 0.9620 | 0.9520 | 0.9610 | 0.9713 | 0.9644 | 0.9679 | 0.9676 | 0.9653 | 0.9560 | 0.9586 | 0.9497 | 0.9771 | 0.9426 | 0.8163 | 0.8925 | 0.9552 | 0.9043 | 1.0000 | |
| | | Sus2_rep3 | 0.9684 | 0.9655 | 0.9728 | 0.9661 | 0.9759 | 0.9722 | 0.9590 | 0.9770 | 0.9837 | 0.9815 | 0.9669 | 0.9713 | 0.9807 | 0.9743 | 0.9734 | 0.9670 | 0.9864 | 0.9566 | 0.8512 | 0.8852 | 0.9471 | 0.9318 | 0.9893 | 1.0000 |

Value
1.0000
0.7185

**S. Fig. 3: Heatmap of Pearson's correlation coefficients display high correlation between biological replicates of Malathion-resistant and -susceptible lines.** Comparisons between lines resistant (Resis#) or susceptible (Sus#) to Malathion when treated with either Malathion at a sublethal dosage or the vehicle control. The numbers following the underscore indicate biological replicates (n=3), each consisting of 8-10 female flies. Comparisons between biological replicates are indicated by black boxes. Highly correlated samples are in red while less correlated samples are in blue.
